# Study of application times gibberellic acid and 2,4-Dichlorophenoxyacetic acid in the plant regeneration from wheat haploid embryos in chromosome elimination method

**DOI:** 10.1101/2020.01.10.902023

**Authors:** Hamed Modirrousta, Raheleh Khademian, Reza Bozorgipour

## Abstract

Wheat is one of the most important cereals, which is very valuable in food. Haploid plants are of particular importance in plant breeding. The wheat seeds produced in the crosses between wheat and maize in the chromosome elimination method without of endosperm and are immature embryo, to prevent the abortions haploid embryos, they must have embryo rescue. Increasing production of haploid plants from produced embryos can improve production efficiency. In this study, With attention their effects gibberellic acid and 2,4-Dichlorophenoxyacetic acid on growth, cell size and cell division, Their use in the production of wheat haploid plant were studied. There was a significant difference at level 1% between the not use and use of gibberellic acid in difference times in the production of haploid from embryos, So that the most haploid plant produced in the use of gibberellic acid in the 4 days after pollination. Also, the use of 2,4-Dichlorophenoxyacetic acid in tiller maintenance liquid culture medium was evaluated at times 48 and 72 hours after pollination. There were a significant difference between these treatments at the 1% level and the most was obtained for wheat haploid plant production with application of 2,4-Dichlorophenoxyacetic acid treatment for 72 hours.

**Highlight:** production of haploid plants plays an important role in wheat breeding. This technique is done to get doubled haploid and absolute homozygous plants in a very short duration of time.

## Introduction

Nowadays the main priorities of wheat breeding are related whit create resistance to disease organisms, changing climate patterns, which need a rapid genetic improvement. The advent and deployment of wheat via maize system of doubled haploid production over the last few decades are discussed as an important option for accelerated wheat breeding. The lack of acute genotypic specificity favours the application of this method in wheat breeding (Srivastava and Bains, 2018). In traditional wheat breeding, the uniformity of lines derived from a breeding population is obtained by repeated selfing which takes several generations to reach homozygosity in loci controlling traits of interest. Using doubled haploid technology, it is possible to attain 100% homozygosity at all loci in a single generation and completely homogeneous breeding lines can be obtained in 1-2 years. Doubled haploid method may significantly reduce cultivar access time. Two major methods for producing wheat doubled haploids are androgenesis and embryo culture using wheat via maize hybridization, which embryo culture being the most effective and widely used method. The method of interspecific hybridization between wheat and maize is laborious but is widely successful for quickly obtaining homozygous lines (Santra et al., 2017; khan et al., 2012). The first reported of the existence haploid plant in nature was reported by Blakeselee et al. (1992) in the *Datura stramorium*. The first haploid plants produced by chromosome elimination method by Kasha and Kao (1970) with use the crosses of *Hordeum vulgare* × *H. bulbosum* reported (Kasha and Kao, 1970). For the first time production of wheat haploid were reported by Laurie and Bennett (1988) using crosses of wheat with maize. Gibberellic acid hormone plays a role in plant growth and development (Schwechheimer, 2008; Spielmeyer et al., 2002). It is also effective in embryo growth and germination (Holdsworth et al., 2008; Yano et al., 2009). In general, during the growth and maturation wheat seed the level of gibberellic acid increase occurs in 15-20 days after pollination (McWha, 1975). Abscisic acid and gibberellic acid affect various aspects of seed development affected, seed dormancy and seed germination are affected by them (Gerjets et al., 2010; Yang et al., 2004). Gibberellic acid plays basic role in the seeds physiological germination and physiological dormancy (Kim and Park, 2008). The use of gibberellic acid externally improves seeds germination (Schopfer et al., 2001). Application of gibberellic acid in wheat increases germination (Tavakol afshari et al., 2011). In all artificial methods of wheat haploid production, it is essential to use the 2,4-Dichlorophenoxyacetic acid. By using the appropriate concentration of this hormone, the amount of haploid production increased up to ten times (Marshall and Molnar-long, 1983). The 2,4-Dichlorophenoxyacetic acid hormone increases the growth pollen tube at the crosses of wheat with maize, which improves this cross and increases the number of seeds and embryos (Wedzony and Vanlammeren, 1996). The use from growth hormone effects on percentage embryo, growth and size of the embryo (Knox, et al., 2000). A simple but effective combination of sugar and sulfurous acid has been introduced to maintenance wheat tillers (Kato and Hayashi, 1985). For embryo rescue, suggested the different times, these times are influenced by genotype and environmental conditions. In crosses wheat with maize, 11-20 days after pollination, for embryo rescue of haploid wheat has been suggested. An early embryo rescue can cause the embryo to not grow properly and stay small and can lose germination in the medium. Of course, the delay in embryo rescue can also cause the embryo to disappear and wrinkle (Sadi et al., 1998). Of pollinated florets, an average of 4.7% of the haploid plant is obtained (Kisana and Nkongolo, 1993). This research was conducted to investigate the effects of times of use of hormones and the remaining residual effect of hormones on improving the production of haploid from wheat embryos produced by chromosome elimination in the wheat crosses with maize.

## Materials and methods

In this research, we have used hybrid F_2_ genotype as the female parent which obtained from Iranian hexaploid bread wheat. Wheat and maize plants were cultivated in the greenhouse. In vegetative and reproductive stages at 20°C and 8 hours darkness and 16 hours lighting were maintenance. After that wheat spikes were coming out from the flag leaf, Florets of spikes were visited continuously. If the stigma were in good condition and the anther did not reach a stage where pollination would occur and were green, Harvested and pollinated in a laboratory with maize pollen without castration was done before the self-pollinated. Immediately after pollination, pollinated spikes were covered with plastic bags for 24 hours. Spikes in a liquid culture medium contains 40 gr sucrose and 8 ml of sulfuric acid and 100 mg of 2,4-Dichlorophenoxyacetic acid per liter, they were maintenance for 48 and 72 hours. After this time, spikes were maintenance in a liquid culture medium containing all of these materials, except for the 2,4-Dichlorophenoxyacetic acid, until the seed harvest was carried out. These tillers were maintenance in a germinator with a humidity content of 65% and darkness for 8 hours and of 16 hours lighting at 20°C. 700 mg of gibberellic acid was dissolved in one liter of water and in times of 2 hours, 2 days, 4 days, 6 days, 8 days and 10 days after pollination were sprayed over the spikes pollinated with pollen. After 18 to 20 days, the seeds were harvested on spikes and they were kept at 4°C for 24 hours. Embryos were isolated from the seeds and the embryos were cultured on a complete MS medium containing 20 gr of sucrose and 8 gr of agar. The embryos were cultured in a separate tube. They were maintenance in incubator at 20°C, after the germination of embryos they are for plant production and to germinator was transferred under the same conditions of tillers maintenance. The number of pollinated florets, number of seeds, number of embryos and number of plants obtained were recorded. Statistical comparisons were performed using Chi-square test. The percentage of seed formation, percentage of embryo formation and percentage plant production using following relationships were obtained, which represents the output frequency.

Percentage of seed formed = Number of seed formed / Number of pollinated florets × 100
Percentage of embryo formed = Number of embryo formed / Number of seed formed × 100
Percentage of plant produced = Number of plant production / Number of embryo formed × 100

## Results

### Effect of 2,4-Dichlorophenoxyacetic acid

Comparison of chi square in between treatments 48 and 72 hours use of 2,4-Dichlorophenoxyacetic acid showed a significant difference at 1% level in the production of haploid plant from the embryos. According to Table 2, the most haploid plant was obtained from embryos, when the tillers maintenance for 72 hours in the liquid culture medium containing 2,4-Dichlorophenoxyacetic acid after pollination. Formation of embryo and production of haploid plants are affected by different concentrations of the 2,4-Dichlorophenoxyacetic acid hormon (Yeshwant and Dilma, 2000). The concentration of 100 mg of 2,4-Dichlorophenoxyacetic acid mg/L is best concentration, because the production of the haploid plant from the embryos will be affected its further increase is negative affected (khan et al., 2012). According to this study, the maintenance of tillers in liquid culture medium contains 100 mg/L of 2,4-Dichlorophenoxyacetic acid mg for 72 hours can significantly increase the production of haploid plant from the embryos formed. In a study by Noga et al. (2016) They investigated the germination of the *Avena sativa* haploid embryos obtained in crosses with maize, they reported the efficiency is germinate embryo of oat to stage of embryo growth and development and hormones used in the medium regeneration of embryos also depend (Noga et al., 2016). Auxins significantly affect the ability of germinate and consequently the production of haploid plant and doubled haploid plant (Marzena et al., 2015). In another study by Dobre and Giura (2015) they reported the examined two hormone methods for production haploid in wheat, they observed that with the use of gibberellic acid and 2,4-Dichlorophenoxyacetic acid, the regeneration of embryos has increased (Dobre and Giura, 2015).

### Effect of gibberellic acid

There was a significant difference at level 1% between not use and use of gibberellic acid in different times in the production of haploid from embryos. So that was produced maximum haploid plant in use of gibberellic acid hormone 4 days after pollination (Table 1). Sitch and Snape (1987) reported that the use of gibberellic acid in the first 10 minute after pollination at the crosses of wheat with wild barley improves, which coincides with the growth of the pollen tube in the stigma prior to influence into the ovary increase the growth speed and embryo development and increase the production efficiency of the haploid from the embryos. Given that hormone gibberellic acid on cell growth and cell division plays a role, this increase in plant production seems to be due to the growth and development of the embryos and organogenesis them. Moieni and Sarrafi (1996) reported that using gibberellic acid in medium the hexaploid wheat anther culture have successful to increase the frequency of haploid production (Moieni and Sarrafi, 1996). The results of this study are correspond with the results of Usha and Khanna (2017) they reported that combined application of gibberellic acid and 2,4-Dichlorophenoxyacetic acid can improve seed production, embryo formation, embryo germination and the regeneration frequency of haploid plant in wheat and maize crosses. This in study reported haploid plant regeneration frequency 36.51%. They also observed that the pattern of embryo growth is different between the same nutrient, due to the effects remaining of the growth regulators (Usha and Khanna, 2017). Laurie and Bennett (1988) reported using gibberellic acid with a concentration of 75 mg/L in their research and using it spray on spikes one day after pollination, growth and embryo formation with gibberellic acid has not improved, low frequency of embryo formation is a show that the vast majority of abortion before development (Laurie and Bennett, 1988). Of course, they used gibberellic acid only in order to produce embryo in spikes wheat in the wheat with maize crosses. The application of gibberellic acid with a concentration of 50 mg/L along with auxin, improves seed and embryo production and embryo development (Matzk, 1991).

**Table 1.** Comparison times use of 2,4-Dichlorophenoxyacetic acid treatments in tillers maintenance medium on haploid plants production from embryos in wheat genotypes

| 2,4-Dichlorophenoxyacetic acid treatments | Number of embryo | Number of plant formation | $\chi^2$ |
| --- | --- | --- | --- |
| 48 hours after pollination | 462 | 167 | 7.08 |
| 72 hours after pollination | 155 | 89 | 13.29 |
| Total | 617 | 256 | 20.37** |

\*\*: significant at 1% level Degrees Freedom: 10

**Table 2.** Comparison of gibberellic acid treatments on haploid plants production in wheat genotypes

| **Gibberellic acid treatments** | **Number of embryo** | **Number of plant formation** | **$\chi^2$** |
| --- | --- | --- | --- |
| No GA<sub>3</sub> | 829 | 192 | 14.65 |
| 2 hours after pollination | 181 | 40 | 7.40 |
| 2 day after pollination | 387 | 138 | 15.45 |
| 4 day after pollination | 64 | 35 | 6.06 |
| 6 day after pollination | 237 | 82 | 7.14 |
| 8 day after pollination | 162 | 45 | 2.96 |
| 10 day after pollination | 104 | 21 | 66.43 |
| Total | 1964 | 553 | 120.09** |

\*\*: significant at 1% level Degrees Freedom: 30

In this study, as shown in Figures 1 and 2 the percentage of seeds and embryos in some treatments was higher than that treatments that produced maximum haploid plant percentage. It should be noted that although the number of seed produced is high but the number of embryos is low or the number of embryos is high but low haploid plants are produced. Increases workload and production efficiency is low. Therefore, the best treatment should produce the most seed and embryo, and finally the haploid plant.

**Fig. 1.**
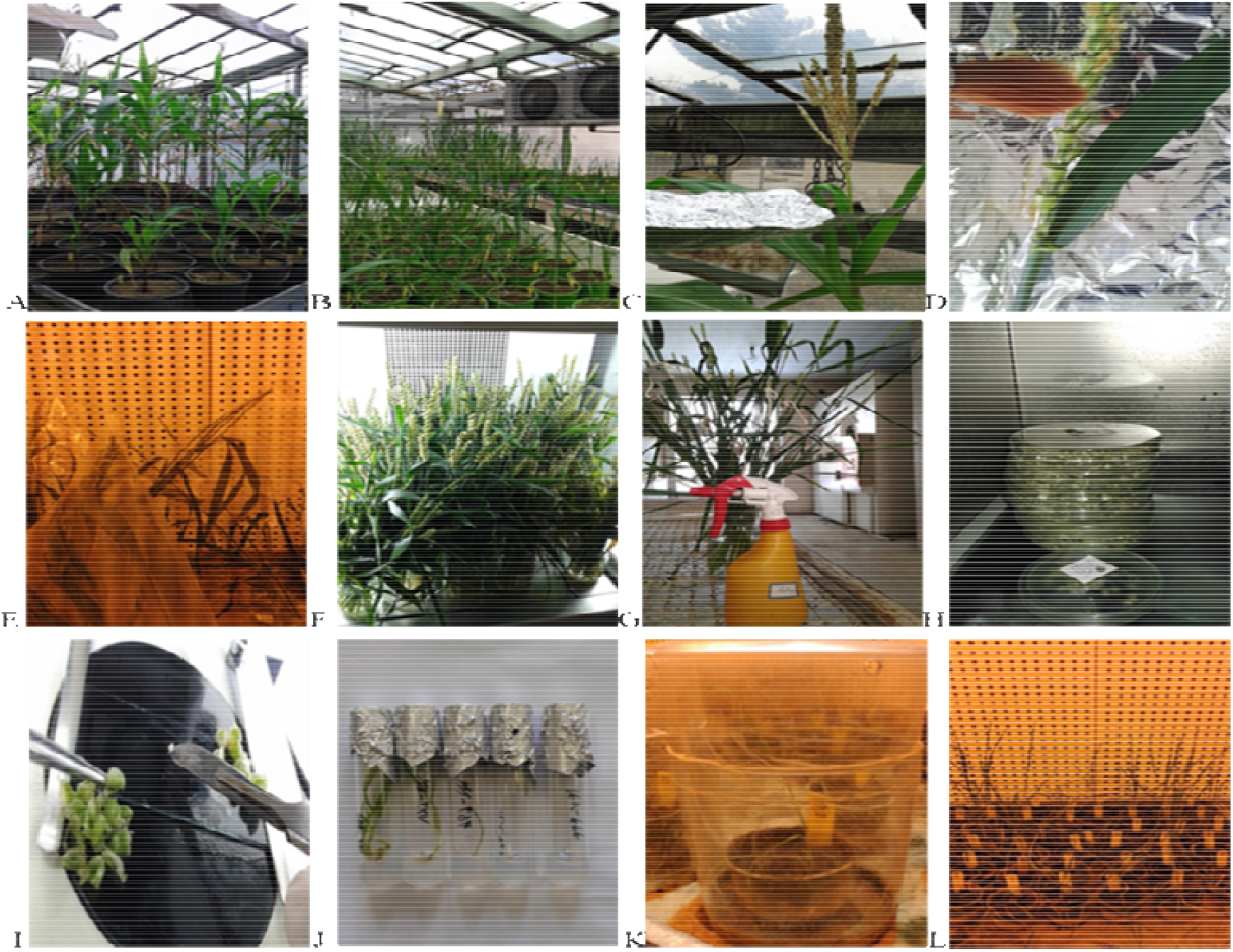
Haploid production process, A: Cultivation of maize plants B: Cultivation of wheat plants C: Collect polle from maize plants D: Pollination of florets of wheat E: Cover the wheat spikes after pollination F: Maintaining the tillers in a liquid culture medium in the germinator G: Spray hormone gibberellic acid on spikes H: The harveste seeds are keep in the refrigerator I: Rescue embryo from seed J: The stages of embryo germination and the production of haploid plant K: The haploid plant cultivate in the soil bed and adaptation L: Haploid plants produce in the tillering stage

**Fig. 2.**
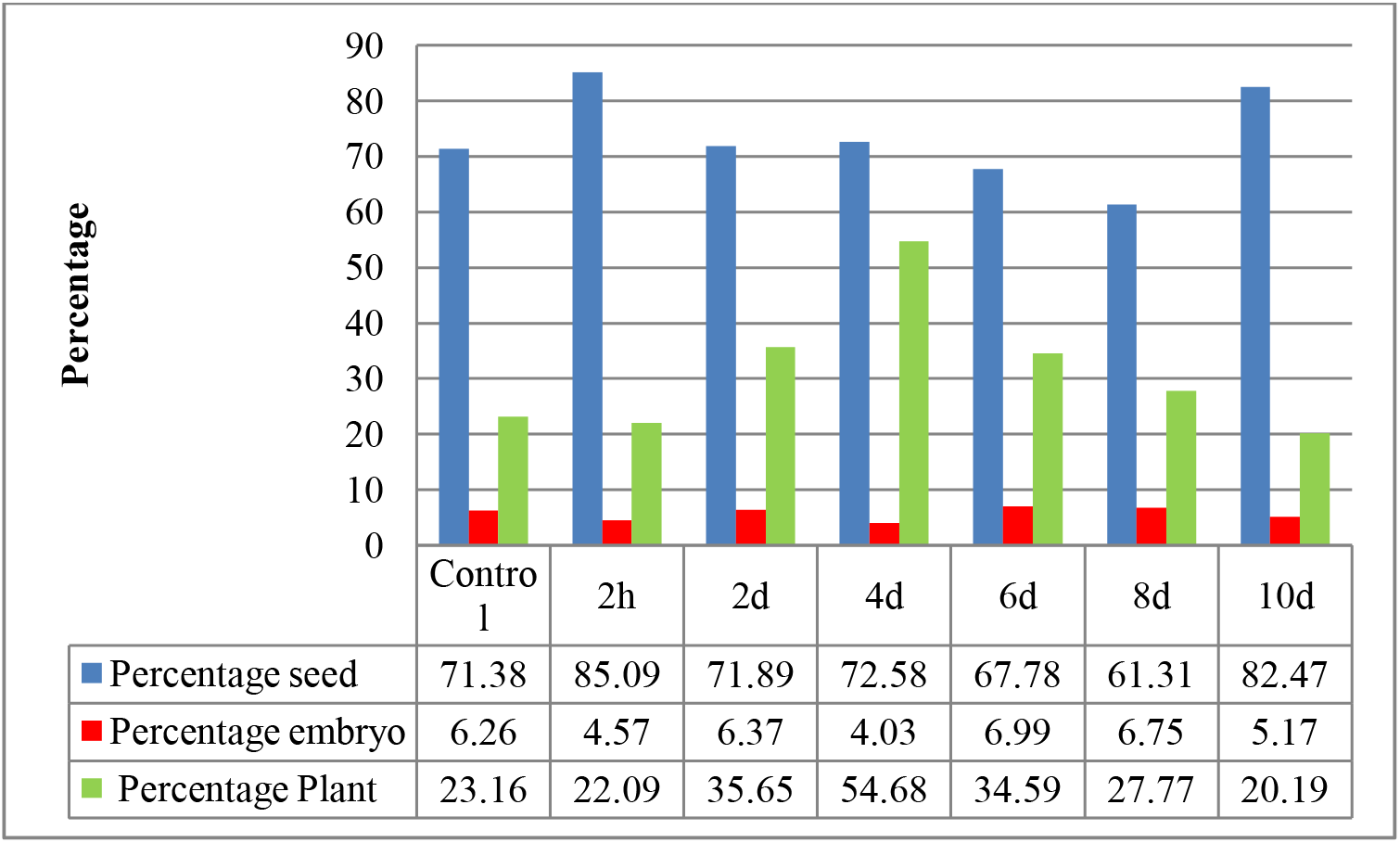
The process of production haploid plants from haploid embryos at different times using gibberellic acid.

**Fig. 3.**
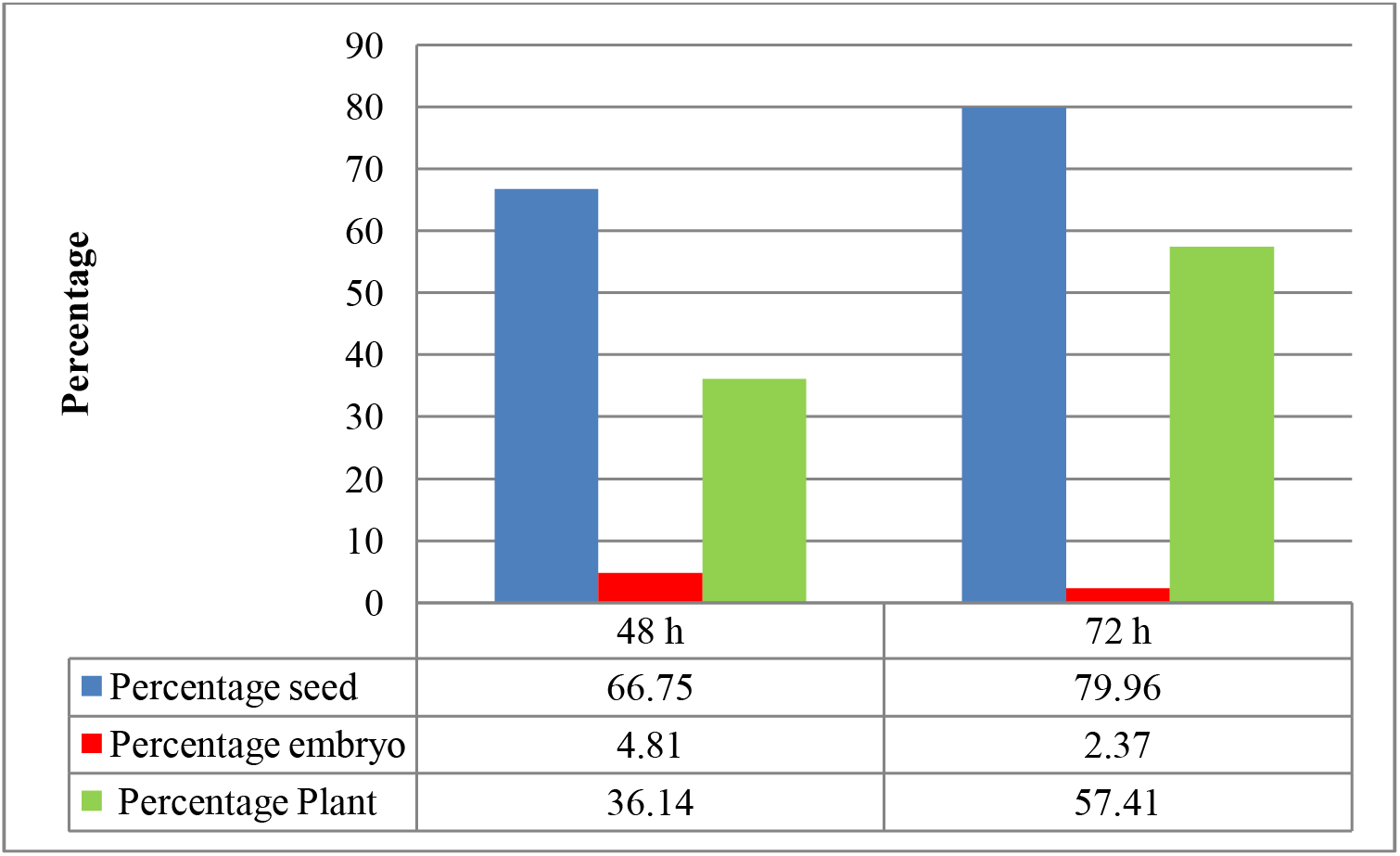
The process of production haploid plants from haploid embryos at different times using 2,4-Dichlorophenoxyacetic acid

### Conclusion

The increase of seed and the haploid embryo when are useful used that increase efficiency the production of haploid plant. Due to the information and experiences obtained from this study, the wheat tiller maintenance for 72 hours in liquid culture medium contains 2,4-Dichlorophenoxyacetic acid and gibberellic acid spray 4 days after pollination recommended for haploid production in bread wheat. Use this method, the wheat haploid plant production improvement in wheat and maize hybridization.

## Acknowledgements

From Research, Seed and Plant Improvement Institute, Agricultural Research, Education and Extension Organization Karaj, Iran, and Imam Khomeini International University, Qazvin, Iran appreciate for providing financial and laboratory facilities for this project. Reza bozorgi pour: Preparation of plant materials, equipment and facilities; Raheleh Khademian: Financial supports.

All authors read and approved the manuscript and the authors declare that they have no conflict of interest.

## References

Afshari RT, Badri S, Abbasi A. 2011. Effects of Gibberellin and Abscisic Acid on Germination, Dormancy Induction as well as Acid and Alkalin Phosphatase Activity in Seed Embryo of Bread Wheat Cultivar, RL4137. Iranian Journal of Crop Sciences 40, 781–789.

Blakese Lee A, Belling J, Farnham M, Bergner A. 1922. A haploid mutant in the jimson weed Datura stramonium. Science 55, 646–647.

Gerjets T, Scholefield D, Foulkes MJ, Lenton JR, Holdsworth MJ. 2010. An analysis of dormancy, ABA responsiveness, after-ripening and pre-harvest sprouting in hexaploid wheat (and pre-harvest sprouting in hexaploid wheat (*Triticum aestivum* L.) caryopses. Journal of Experimental Botany 61, 597–607.

Dobre S, Giura A. 2015. Aspects regarding wheat genotypic response to PGR treatments applied in vivo. Journal of Horticulture, Forestry and Biotechnology 19, 21–24.

Holdsworth MJ, Bentsink L, Soppe WJ. 2008. Molecular networks regulating Arabidopsis seed maturation, after-ripening, dormancy and germination. The New phytologist 179, 33–54.

Kasha K, Kao K. 1970. High frequency haploid production in barley (*Hordeum vulgare* L.). Nature 225, 874–876.

Kato K, Hayashi H. 1985. Modification method to obtain mature dry seeds by sucrose solution culture of detached wheat ears. Agric Sci 33, 63–70.

Khan MA, Shaukat S, Ahmad J, Kashif M, Khan AS, Iqbal Z. 2012. Use of intergeneric cross for production of doubled haploid (*Triticum aestivum*). Sci., Tech. and Dev 31, 295–300.

Kim SG, Park CM. 2008. Gibberellic acid-mediated salt signaling in seed germination. Plant Signal Behav 3, 877–879.

Kisana NS, Nkongolo KK. 1993. Production of doubled haploids by anther culture & wheat × maize method in a wheat breeding program. plant breeding 110, 96–102.

Knox RE, Clark JM, Depauw RM. 2000. Dicamba and growth condition effects on doubled haploid production in durum wheat crossed with maize. Plant Breeding 119, 289–298.

Laurie DA, Bennett MD. 1988. The production of haploid wheat plants from wheat × maize crosses. Theor. Appl. Genet 79, 393–397.

Marshall D, Molnar-long M. 1983. Effect of 2,4-D on partenocarpy and cross compatibility in wheat. Cellular & Molecular Biology Letters 6, 313–318.

Matzk F. 1991. A novel approach to differentiated embryos in the absence of endosperm. Sexual Plant Reproduction 4, 88–94.

McWha JA. 1975. Changes in abscisic acid levels in developing grains of wheat (*Triticum aestivum* L). J. Exp. Botany 26, 823–827.

Moieni A, Sarrafi A. 1996. The effects of gibberllic acid, phenylethylamine, 2,4-D, and genotype on androgenesis in hexaploid wheat (*Triticum aestivum*). Cereal Research Communications 24, 139–145.

Sadi N, Chylah O, Chlyah H. 1998 Production of green haploid durum wheat plant by pollination of wheat with maize. Can. J. Bot 76, 652–656.

Santra M, Wang H, Seifert S, Haley S. 2017. Doubled Haploid Laboratory Protocol for Wheat Using Wheat-Maize Wide Hybridization. In: Bhalla P, Singh M, eds. Wheat Biotechnology: Methods in Molecular Biology. Humana Press: New York, 235–249.

Schopfer P, Plachy C, Frahry G. 2001. Release of reactive oxygen intermediates (superoxide radicals, hydrogen peroxide, and hydroxyl radicals) and peroxidase in germinating radish seeds controlled by light, gibberellin, and abscisic acid. Plant Physiol. 125, 1591–1602.

Schwechheimer C. 2008. Understanding gibberellic acid signaling - are we there yet?. Current Opinion in Plant Biology 11, 9–15.

Sitch LA, Snape jw. 1987. Factors affecting haploid production in wheat using the *Hordeum bulbosum* system. 1. Genotypic and environmental effects on pollen grain germination, pollen tube growth and the frequency of fertilization. Euphytica 36, 483–496.

Srivastava P, Bains NS. 2018. Accelerated Wheat Breeding: Doubled Haploids and Rapid Generation Advance. In: Gosal S, Wani S eds. Biotechnologies of Crop Improvement. Cham: Springer, 437–461.

Usha, P., Khanna VK. 2017. Effect of hormonal treatments on haploid formation and in vitro haploid regeneration in wheat × maize system. International Journal of Plant Sciences 12, 234–239.

Warchoł M, Skrzypek E, Nowakowska A, Marcińska I, Czyczyło-Mysza I, Dziurka K, Juzoń K, Cyganek K. 2016. The effect of auxin and genotype on the production of *Avena sativa* L. doubled haploid lines. Plant Growth Regulation 78, 155–165.

Warchoł M, Czyczyło‑Mysza I, Marcińska I, Dziurka K, Noga A, Skrzypek E. 2018. The effect of genotype, media composition, pH and sugar concentrations on oat (*Avena sativa* L.) doubled haploid production through oat × maize crosses. Acta Physiologiae Plantarum 40:93.

Wedzony M, Vanlammeren A. 1996. Pollen Tube Growth and Early Embryogenesis in Wheat × Maize Crosses Influenced by 2,4-D. Annals of Botany. 77, 639–647.

Yano R, Kanno Y, Jikumaru Y, Nakabayashi K, Kamiya Y, Nambara E. 2009. CHOTTO1, a putative double APETALA2 repeat transcription factor, is involved in abscisic acid-mediated repression of gibberellin biosynthesis during seed germination in Arabidopsis. Plant physiology 151, 641–654

Yang J, Zhang J, Wang Z, Xu G, Zhu Q. 2004. Activities of key enzymes in sucrose-to-starch conversion in wheat grains subjected to water deficit during grain filling. Plant Physiol 135, 1621–1629.

Yeshwant RM, Dilma CA. 2000. Somaclonal variation for disease resistance in wheat and production of dihaploids through wheat × maize hybrids. Genetics and Molecular Biology 23, 617–622.

Zhang G, Friebe B, Raupp G, Harrison S, Gill B. 1996. Wheat embryogenesis and haploid production in wheat × maize hybrids. Ephytica 90, 315–324.

